# Cell-free expression of Nipah virus transmembrane proteins for proteoliposome vaccine design

**DOI:** 10.1101/2024.07.26.605347

**Authors:** Vivian T. Hu, Shahrzad Ezzatpour, Ekaterina Selivanovitch, Jordan Carter, Julie Sahler, Richard Ayomide Adeleke, Avery August, Hector C. Aguilar, Susan Daniel, Neha P. Kamat

**Affiliations:** Department of Biomedical Engineering, Northwestern University, Evanston, IL, 60208 USA; Center for Synthetic Biology, Northwestern University, Evanston, IL, 60208 USA; Department of Microbiology and Immunology, College of Veterinary Medicine, Cornell University, Ithaca, NY 14853, USA; Robert Frederick Smith School of Chemical and Biomolecular Engineering, Cornell University, Ithaca, NY 14853, USA

**Keywords:** cell-free protein synthesis, liposomes, vaccine, Nipah virus, transmembrane proteins

## Abstract

Membrane proteins expressed on the surface of enveloped viruses are potent antigens in a vaccine, yet are difficult to produce and present due to their instability without a lipid scaffold. Current vaccination strategies that incorporate viral membrane proteins, such as live attenuated viruses, inactivated viruses, or extracellular vesicles, have limitations including lengthy production time, poor immunogenicity, extensive processing steps, and/or poor stability. Cell-free protein synthesis of viral membrane proteins offers a rapid, one-step method to assemble vaccine nanoparticles via cotranslational folding of membrane proteins into nanoscale liposomes. Here, we develop a vaccine candidate for the deadly Nipah virus (NiV), a highly lethal virus listed by the World Health Organization as a priority pathogen, by cell-free expressing two full-length Nipah virus membrane proteins. We demonstrate that both NiV fusion protein (NiV F) and NiV glycoprotein (NiV G) can be expressed and cotranslationally integrated into liposomes and that they fold into their native conformation. We find the removal of a signal peptide sequence and alteration of liposome lipid composition improves viral membrane protein incorporation. Furthermore, a lipid adjuvant, monophosphoryl lipid A (MPLA), can be readily added to liposomes without disrupting protein-vesicle loading or protein folding conformations. Finally, we demonstrate that our generated liposomal formulations lead to enhanced humoral responses in mice compared to empty and single-protein controls. This work establishes a platform to quickly assemble and present membrane antigens as multivalent vaccines that will enable a rapid response to the broad range of emerging pathogenic threats.

## Introduction

Viral membrane proteins are an important target for the immune system. Accessible on the surface of viral particles, these membrane proteins are critical in receptor binding and viral entry. Accordingly, neutralizing antibodies produced against epitopes on the surface of these proteins can confer strong protection in an infected host (1). Several features of viral membrane proteins have been shown to be important for their immunogenicity including their orientation (2), conformation (3, 4), stability (5, 6), and surface presentation (7). In particular, the nonpolar transmembrane domain of these proteins appears to help present viral antigens in a more native-like conformation, and thus are likely to be important to include (8-13). However, incorporating membrane proteins into vaccine formulations is challenging due to their hydrophobic nature and instability without a lipid membrane. To address this constraint, viral membrane proteins that are expressed with their transmembrane domains are often assembled as components of extracellular vesicles (14, 15) or virus-like particles (VLPs). These approaches require that the proteins are genetically encoded in plasmids in host cells, produced by that cell, and later extracted in the form of vesicles or VLPs from the cell. These methods require extensive steps in the way of cell culture, extraction, and purification and new biomanufacturing strategies to present viral, membrane-protein antigens in a biomimetic manner are needed.

Cell-free protein synthesis (CFPS) systems, which contain extracted protein machinery from cells, are a promising tool for vaccine development because they can rapidly produce clinically-relevant amounts of protein, can be freeze-dried for distribution, and prevent the need for cold chain storage (16-19). Cell-free expressed (CFE) membrane proteins can be rapidly added to the surface of or cotranslationally integrated into amphiphilic scaffolds like liposomes (20-23). This strategy results in one-step assembly of lipid nanoparticles with membrane-integrated proteins, preventing the need for multi-step chemical conjugation or protein reconstitution schemes (23-25). While this technique has largely been explored in fundamental membrane protein folding and structure studies (20, 26-28) and synthetic cell work (21, 22), employing this assembly strategy to design vaccines and therapeutics is an emerging and promising approach (16).

While many amphiphilic scaffolds can be used to incorporate and present viral membrane proteins, bilayer-based scaffolds, such as liposomes, are particularly useful carriers for nanoparticle vaccines because they can recapitulate many features of the virus membrane envelope. Features such as particle size, charge, fluidity, and phospholipid makeup can be tuned to influence vaccine activity (29-34). In addition, simply attaching antigenic proteins to the outside of a scaffold has been shown to improve uptake by antigen-presenting cells and lymph node retention, overall producing stronger and lasting neutralizing antibody titers and protection (7, 35). Further, nanoparticles containing hydrophobic bilayers are amenable to lipid adjuvants and antigens that can be colocalized with the other vaccine components (36-39) and can accommodate multiple membrane proteins for the design of multivalent or universal vaccines. Finally, these scaffolds can be designed in a way that allows attached or integrated proteins to be spatially patterned (40). Taken together, using CFPS to decorate bilayer-based nanoparticles with viral membrane proteins is a promising strategy to create rapidly produced, low cost, and shelf-stable vaccines with the potential for enhanced potency relative to current strategies (Fig. 1A).

**Fig. 1.**
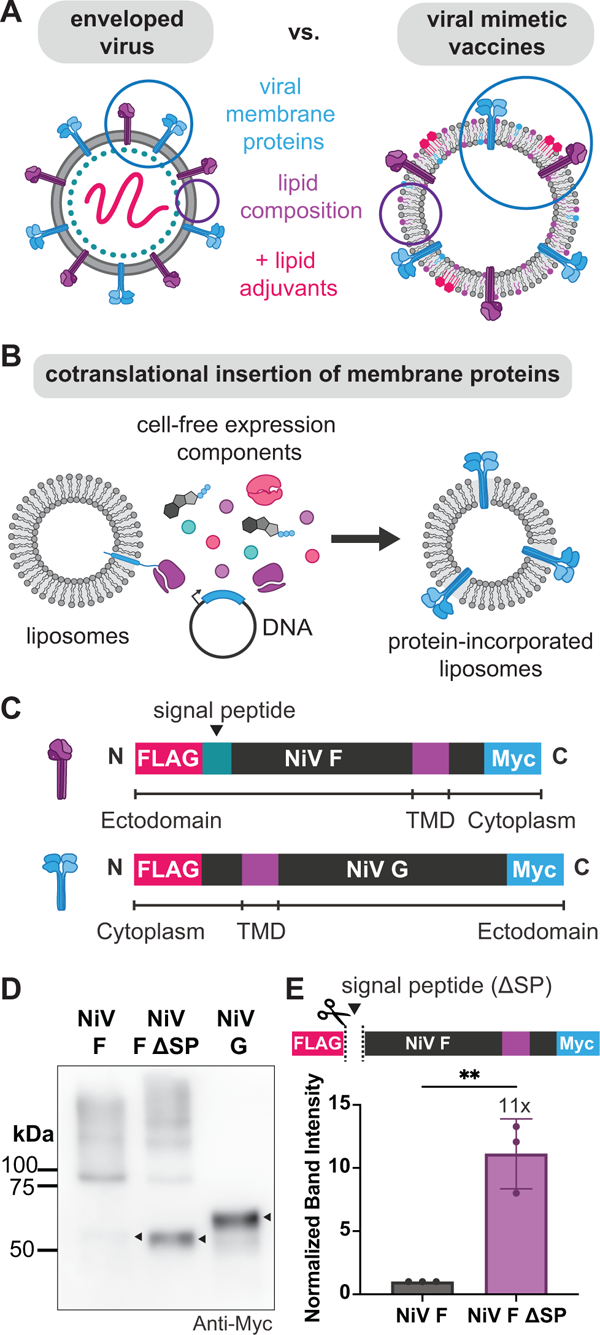
Assembly of viral mimetic particles via cotranslational integration of viral membrane proteins into liposomes. (**A**,**B**) Schematic of experimental design: **(A)** Liposomes can be used as scaffolds to present viral transmembrane proteins for use as viral mimetic particles **(B)** Schematic workflow of viral mimetic vesicle preparation in which cell-free protein synthesis systems are used to produce viral membrane proteins. Supplementing reactions with hydrophobic supplements such as liposomes leads to cotranslational integration of the synthesized proteins. **(C)** Design of genes for two single-pass transmembrane proteins: Nipah virus fusion (NiV F) and Nipah virus attachment (NiV G). **(D)** Western blot analysis of purified DOPC liposomes after cell-free expression reactions reveals full-length proteins NiV F, NiV F ΔSP, and NiV F proteins are present and membrane associated. Antibodies against a C-terminal Myc tag were used for blot analysis. **(D)** Deletion of the signal peptide sequence in Type I transmembrane protein NiV F (NiV F ΔSP) enhances NiV F expression and vesicle association by roughly 11-fold. Error bars represent the standard deviation for n = 3 independent replicates. \*\**P*≤ 0.01; *P* value is from a two-tailed t-test.

In this study, we used an *in vitro* CFPS system to synthesize and integrate full-length viral membrane proteins from Nipah virus (NiV), a neurological and respiratory disease with a high case fatality rate of 40 to 75% (Fig. 1B) (41), into nanoscale liposomes. Because there are no approved therapies or vaccines for this disease, the World Health Organization (WHO) has identified NiV as a priority pathogen for research and development (42), making NiV a critical target for vaccine design. We investigated how changing lipid composition impacts cotranslational expression of NiV F and G proteins and their co-incorporation into liposome membranes. We also assessed the conformation of the resulting proteins using conformational antibodies. Lastly, we performed mice vaccination studies with our assembled nanoparticle vaccines to assess neutralizing antibody production and cellular responses. Together, this work describes a platform for the design of cell-free modified nanoparticle vaccines that is applicable to a range of viral diseases, and in which several features of the vaccine (protein sequence and identity, lipid adjuvants, encapsulated cargo) can be modularly altered toward improved, rapid generation of effective vaccines.

## Results

### Viral Membrane Proteins Nipah Fusion and Nipah Attachment Can Be Expressed Using Cell-Free Protein Synthesis into Liposomes

We assembled nanoscale liposomes containing two cell-free expressed viral membrane proteins from NiV. NiV contains two immunogenic, single-pass transmembrane glycoprotein antigens required for viral entry into host cells, the receptor binding glycoprotein (NiV G), which binds host cell receptors ephrinB2 and ephrinB3, and the fusion protein (NiV F), which mediates fusion of the viral and cellular membranes during viral entry into host cells (43-49). Accordingly, antibodies produced against these proteins can combat viral infections by preventing enveloped viruses from binding and fusing to target host cells as well as activating cells in the innate and adaptive immune system. For each transmembrane protein, we designed constructs with an N-terminal FLAG and C-terminal Myc peptide tags to aid in western blot analysis and orientation studies (Fig. 1C). The viral protein sequences were optimized for use in bacterial cell-free protein synthesis systems, which are more cost-effective, optimizable, scalable, and widely used compared with eukaryotic systems (50-52).

We first confirmed that NiV F and G proteins could separately be expressed and integrated in liposomes. Unilamellar, nanoscale liposomes were assembled from 1,2-dioleoyl-sn-glycero-3-phosphocholine (PC) lipids via thin film hydration techniques and extruded to 100 nm in average size. The liposomes also contained 1% of a biotinylated lipid to enable liposome purification away from cell-free expression components in later steps. NiV F and G were expressed individually in the presence of liposomes. To isolate the expressed protein that cotranslationally associated with liposomes from soluble and aggregated protein in solution, we developed a purification scheme using magnetic streptavidin beads that bind to biotinylated liposomes (Fig. S1). As shown in Fig. 1D, full-length NiV F (∼63 kDa, lane 1) and NiV G (∼69 kDa, lane 3) were expressed and associated with purified liposomes.

### Alterations in the Nascent Peptide Sequence of Nipah Transmembrane Proteins and Liposome Composition Can Improve Protein Expression and Association with Liposomes

The amount of protein present on a viral mimetic particle is crucial for vaccine efficacy because the particle should contain sufficient amounts of viral protein to generate high cellular and humoral immune responses as well as contain enough antigen for pathogen specificity (53, 54). Toward achieving higher levels of membrane protein expression and transmembrane integration into liposomes, we wondered if we could improve NiV F expression by changing the design of its protein sequence. NiV F is a type I transmembrane protein and has a 26 amino acid hydrophobic signal peptide (SP) at its N-terminus preceding an extracellular N-terminus. This peptide is important in cells for membrane targeting and insertion into the secretory pathway (55). However, this peptide is unnecessary in our cell-free system that lacks a signal recognition particle and we hypothesized that the presence of the hydrophobic peptide in the nascent peptide chain might hinder the cotranslational integration of the membrane protein into liposome membranes. Further, we have previously found that sequence alterations in the nascent chain of a cell-free expressed membrane protein can have a profound impact on cell-free membrane protein expression (56). Specifically, we identified that alterations in protein sequence that inhibit protein-lipid interactions lead to membrane proteins that cannot successfully move off the ribosome and insert into membranes. As a result, stalled ribosomes can inhibit subsequent expression by competitively binding other nascent proteins and ribosomes. To potentially improve both cotranslational integration into liposomes and overall protein expression, we deleted the sequence in NiV F encoding the signal peptide (NiV F ΔSP). We expressed NiV F ΔSP in PC liposomes and western blot analysis showed a marked increase in membrane associated protein for the NiV F ΔSP sample as compared to the NiV F sample (Fig. 1D). The band intensities of NiV F and NiV F ΔSP were quantified and normalized to the band intensity of NiV F, revealing roughly an 11-fold increase in membrane protein association due to deletion of the signal peptide (Fig. 1E). To explore the generality of our finding that signal sequence deletion can improve cell-free expression of type I membrane proteins, we explored two other viral membrane proteins: SARS-CoV-2 spike (S) and Influenza hemagglutinin (HA). We expressed these proteins using CFPS and in the presence of PC liposomes and measured a 14-fold increase in membrane associated protein for Spike (S) ΔSP relative to the full-length S protein. In contrast, deletion of the signal peptide in HA had no significant effect on HA association with PC liposomes (Fig. S2). Upon visualizing the molecular structures, we noticed that the signal peptide is in the globular head for NiV F and S yet in the stalk region close to the more amphiphilic transmembrane domains for HA. Thus, we hypothesize that NiV F and S expression were significantly enhanced when this hydrophobic sequence was removed from their soluble head yet did not affect expression of HA presumably because its signal sequence is near a more hydrophobic stretch of the protein. Our results modifying the nascent peptide sequence of the selected viral transmembrane proteins points to an additional handle to improve cell-free expression of type I transmembrane proteins that should be tested for individual membrane proteins of interest. Unlike NiV F, NiV G is a type II transmembrane protein lacking a signal peptide. Due to the improved expression in lipophilic environments, we used the NiV F ΔSP mutant and the full-length NiV G proteins for subsequent studies.

### Changing Membrane Composition Improves Membrane-Association of Cell-free Expressed Proteins

We next explored the capacity of liposome composition to enhance the expression and integration of NiV transmembrane proteins. The cell-free expression of transmembrane proteins has been shown to be highly sensitive to the composition of amphiphilic scaffolds present in the reaction (20-22, 57). We hypothesized that changing the biochemical makeup of liposomes could promote the cotranslational folding of viral membrane proteins into the bilayer, in turn improving the amounts of protein associated with the membrane.

Specifically, we evaluated the capacity of changes in the lipid headgroup to impact protein expression, while keeping the lipid acyl chain length and saturation constant (Fig. 2A). Three compositions served as the basis for investigation of the composition studies: as a control composition, we chose to use 100% 1,2-dioleoyl-sn-glycero-3-phosphocholine (DOPC) liposomes containing a phosphocholine (PC) headgroup which is a neutral zwitterion. Phosphatidylcholine is the most abundant headgroup in mammalian cells and is a common component of liposomes present in cell-free membrane protein expression reactions (22, 58, 59). In addition, we looked at lipid compositions with 20 mol% 1,2-dioleoyl-sn-glycero-3-phosphoethanolamine (DOPE), which we refer to as PC/PE composition. Compared to phosphatidylcholine, phosphatidylethanolamine is also zwitterionic yet has a more conical shape that introduces negative curvature in to a bilayer membrane (24). This negative curvature adds defects in the membrane that can potentially improve the cotranslational insertion of membrane proteins (60). In addition, PC- and PE-containing lipids are the most abundant phospholipids in eukaryotes, with PE comprising 15-25% of lipids in mammalian cells (61). Finally, toward a more biomimetic scaffold for the viral membrane proteins, we looked at a third composition of eukaryotic lipids that included 5 mol% 1,2-dioleoyl-sn-glycero-3-phospho-L-serine (DOPS) in addition to 20 mol% DOPE and 75 mol% DOPC, which we refer to as PC/PE/PS. The phosphatidylserine headgroup is negatively charged and makes up around 10-20% of the phospholipids in the plasma membrane of many cells, from where many enveloped viruses including NiV bud during assembly (62-64). In addition, PS, which is a marker for apoptosis in eukaryotic cells, becomes present on the surface of cells after viral infection (65) and is present on the outer leaflet of virus envelopes (66). The negative charge of PS is also predicted to improve storage stability of the liposomes since they electrostatically repel each other, reducing aggregation (67). While viral membranes also include other components such as sterols, sphingolipids, and glycolipids, we focused on tuning glycerophospholipid compositions via the aforementioned lipid changes since they are the most abundant lipids in the plasma membrane.

**Fig. 2.**
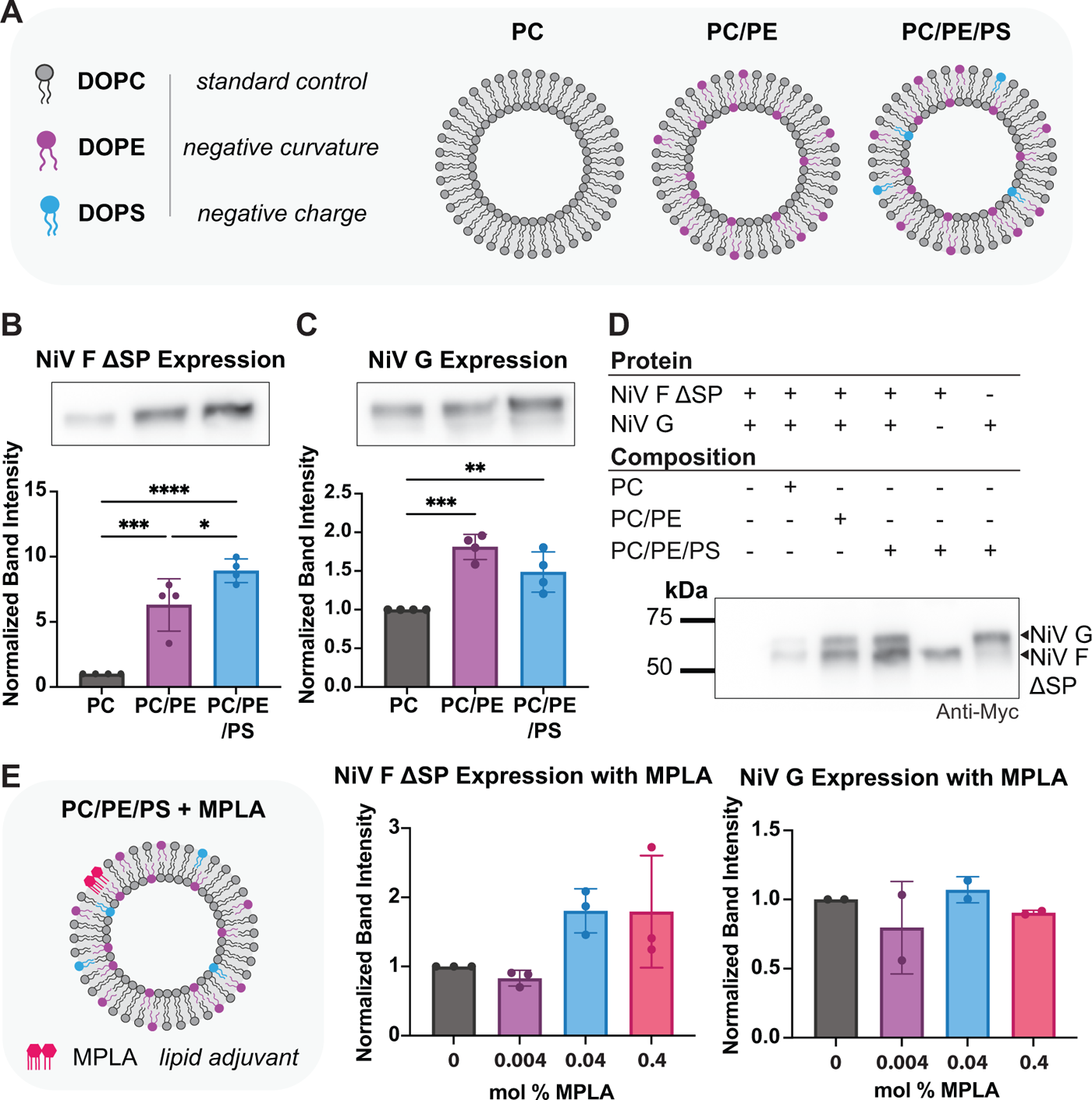
Alteration of liposome membrane composition enhances NiV F and NiV G association with liposome membranes. **(A)** Liposomes were assembled with lipids with different phospholipid headgroups (phosphatidylcholine, phosphoethanolamine (PE), and phosphatidylserine (PS)). Liposomes were either 100% DOPC (PC), 4:1 PC:PE (PC/PE), or 15:4:1 PC:PE:PS (PC/PE/PS), **(B)** NiV F ΔSP expression and integration into liposomes increases by 6.3- and 8.9-fold in the presence of 4:1 PC:PE (PC/PE) and 15:4:1 PC:PE:PS (PC/PE/PS), respectively, with respect to PC liposomes. **(C)** NiV G expression and integration into liposomes increases by 1.8- and 1.5-fold in PC/PE and PC/PE/PS liposomes, respectively, with respect to PC liposomes **(D)** Co-expression of NiV F ΔSP and NiV G in the presence of PC, PC/PE, and PC/PE/PS liposomes results in detection of both proteins associated with liposomes, assessed via western blot analysis. **(E)** Semi-quantitative western blot analysis reveals that inclusion of MPLA lipid at amounts of 0%, 0.004%, 0.04%, and 0.4% in PC/PE/PS liposomes does not significantly affect the subsequent expression and vesicle association of NiV F ΔSP and NiV G. Error bars represent the standard deviation for n = 4 (B,C), n=3 (E middle), or n=2 (E right) independent replicates. \**P* ≤ 0.05; \*\**P* ≤ 0.01; \*\*\**P* ≤ 0.001; \*\*\*\**P* ≤ 0.0001; *P* values are generated using one-way analysis of variance (ANOVA) and Tukey’s multiple comparisons test.

We first expressed NiV F ΔSP in PC, PC/PE, and PC/PE/PS liposomes. Liposomes contained 1% biotinylated lipid (18:1 Biotinyl Cap PE) for streptavidin-mediated purification and 0.1% Cy5.5 fluorescent lipid dye (18:1 Cy5.5 PE). To further assist cell-free folding of the membrane proteins that contain multiple disulfide bonds, we used the PURExpress® Disulfide Bond Enhancer, which helps with oxidation of cysteine thiols, throughout all subsequent *in vitro* studies. For our comparison, we performed a semi-quantitative western blot analysis on liposomes purified after the cell-free reaction. We quantified the band intensities of the protein expressed in the different compositions and normalized the band intensity values to the total fluorescence of the Cy5.5 membrane dye measured of purified vesicles to account for variability during sample purification. Afterwards, these normalized band intensity values were then normalized to this value obtained for the PC liposomes to allow for standardization in subsequent replicates. Following this analysis, we observed that adding DOPE to the PC liposomes significantly improved NiV F ΔSP membrane association by 6.3-fold compared to PC liposomes, and the inclusion of both DOPE and DOPS in the PC/PE/PS liposomes enhanced membrane association by 8.9-fold (Fig. 2B) relative to PC liposomes. We performed the same study to investigate expression of NiV G in the PC, PC/PE, and PC/PE/PS liposomes and saw that compositional changes likewise improved NiV G expression and liposomal association levels compared to the PC group. The changes were significant, albeit to a lesser extent than those seen for NiV F ΔSP, with a 1.8-fold improvement in NiV G expression in PC/PE liposomes and a 1.5-fold increase in PC/PE/PS liposomes (Fig. 2C) relative to PC liposomes. Liposome size was measured after expression via dynamic light scattering (Malvern Zetasizer) (SI Table 1, Figure S3). Finally, toward a multivalent vaccine, we co-expressed both type I NiV F ΔSP and type II NiV G transmembrane proteins in PC, PC/PE, and PC/PE/PS liposomes and performed western blot analysis to confirm their incorporation into purified PC/PE/PS liposomes (Fig. 2D), confirming that both proteins could be co-incorporated into liposome membranes.

We also analyzed the orientation of NiV F ΔSP and NiV G across the compositions via flow cytometry (Fig. S4). Both NiV F ΔSP and NiV G proteins were designed to contain an N-terminal FLAG tag and C-terminal Myc tag. For an orientation assessment, we labeled magnetic Protein A/G beads with PE conjugated anti-FLAG or anti-Myc antibodies. NiV F ΔSP or NiV G were expressed into PC, PC/PE, or PC/PE/PS liposomes containing Cy5.5. Depending on the orientation of single-pass NiV F ΔSP and NiV G in the liposomes, the proteins should bind to the antibody-functionalized bead if the tags are surface-accessible and should not bind if the tags are in the protected luminal part of the liposomes. The beads with bound liposomes were analyzed via flow cytometry. Colocalization of Cy5.5 fluorescence (from membrane dyes) to the beads indicates that the specific protein tag in question, FLAG or Myc tag, was accessible on the outside of the vesicle. While membrane associated protein was detected in both orientations across compositions for NiV F ΔSP and NiV G, we observed significant increases in protein-mediated binding to antibody beads relative to the control empty liposomes when the proteins were in their native orientation. In contrast, protein-mediated binding to antibody beads in the “upside-down” orientation across compositions was nonsignificant relative to empty liposome controls for both NiV F ΔSP and NiV G. Beyond better mimicking the presentation of these proteins in native viruses, our orientation studies also provided initial evidence that the NiV proteins are integrated into bilayers versus passively adsorbed on the liposomal surface. These results were also supported via another independent assay using a broad-spectrum protease cleavage assay with proteinase K (Fig. S5). Here, protein fragments on the outside of the liposome are digested while fragments integrated in the bilayer or inside of the liposome are protected. Samples containing the detergent Triton X-100 were used as a control, because all protein regions should be digested for liposomes that are disrupted by the detergent. Depending on the orientation of the protein, either the FLAG- or Myc-tag should be protected and detectable on via western blot analysis. We observed that compared to respective samples treated with 0.1% Triton, NiV G appears to have predicted banding on a western blot whereas NiV F ΔSP has detectable protein in both orientations.

Next, we wondered if adding a lipid adjuvant to liposomes would affect subsequent expression and association of NiV F ΔSP and NiV G proteins. Specifically, liposomes provide an amphiphilic surface that can incorporate a range of amphiphilic components like other lipids. Of interest to our design was the inclusion of adjuvants, which are commonly used in vaccine formulations to improve their efficacy by activating innate immune responses. One adjuvant that is commonly used in lipid-based systems is monophosphoryl lipid A (MPLA) (68). MPLA is a modified endotoxic lipid that interacts with TLR4 to promote antigen presentation and T-cell activation, and it has been shown to be an effective adjuvant in enhancing Th1 responses, which are important for intracellular infections (36, 69, 70). While it has previously been demonstrated that MPLA does not affect the cell-free expression of *Pseudomonas aeruginosa* outer membrane protein OprF, OprF consists of an eight stranded β-barrel (71), which has a different mechanism of membrane insertion than the α-helical Nipah proteins in our study. We therefore sought to characterize the effect MPLA inclusion in liposomes would have on cell-free expression and cotranslational folding of the NiV F and G proteins.

To assess the effects of MPLA inclusion on NiV F and G protein expression and association, we assembled PC/PE/PS liposomes with varied concentrations of MPLA (0 mol%, 0.004 mol%, 0.04 mol%, and 0.4 mol% MPLA in PC/PE/PS liposomes). We analyzed the resulting amount of NiV F ΔSP and NiV G that were membrane associated after cell-free expression and liposome purification. Semi-quantitative western blot analysis of NiV F ΔSP and NiV G was performed (Fig. 2E) and we normalized the protein band intensities based on vesicle fluorescence from a membrane dye in each run. This quantity is reported as normalized to the 0% MPLA group to allow for standardization across runs. Our results indicate the presence of MPLA did not significantly affect the amounts of NiV F ΔSP or NiV G associated with liposomes across the range of adjuvant tested, likely allowing for further flexibility in tuning the amounts of MPLA per dose based on cell-free reaction yields.

### Cell-free Expressed Viral Membrane Proteins Maintain Native Folding Conformation

We wondered if cell-free expressed and membrane associated NiV F ΔSP and NiV G proteins retain their expected conformation. In peptide-based nanoparticle vaccines, integrated protein orientation and conformation are particularly critical for eliciting neutralizing antibody responses against the native virus (54, 72). However, since bacterial cell-free reactions occur outside of the cell, they commonly lack the chaperones, translocons, and folding enzymes that assist in protein folding (60). Improper folding can lead to aggregation in the reaction as well as a potentially less robust immune response.

To assess the extent to which CFE proteins folded in their native conformation, we used conformational antibodies developed against NiV F and NiV G proteins (Fig. 3A) (73). For this study, NiV F ΔSP and NiV G were individually expressed in PC/PE/PS liposomes containing 0.1% Cy5.5. Liposomes were then incubated individually with separate monoclonal antibody (Mab)-functionalized beads that bind conformation-specific epitopes on NiV F or NiV G. An empty liposome control with no protein template was used to measure background fluorescence due to nonspecific vesicle adsorption to antibody beads. The conformational antibodies used were previously described as having a strong signal in flow cytometry and extremely weak to no signals in denaturing western blots (74-77). In our assay, we expected that protein that is surface accessible on the liposome will bind to the antibody. Flow cytometry should then detect the colocalization of beads and liposome membrane fluorescence, with colocalization of liposome fluorescence and beads indicating that proteins in the liposome adopt a native conformation and are binding to the antibody beads.

**Fig. 3.**
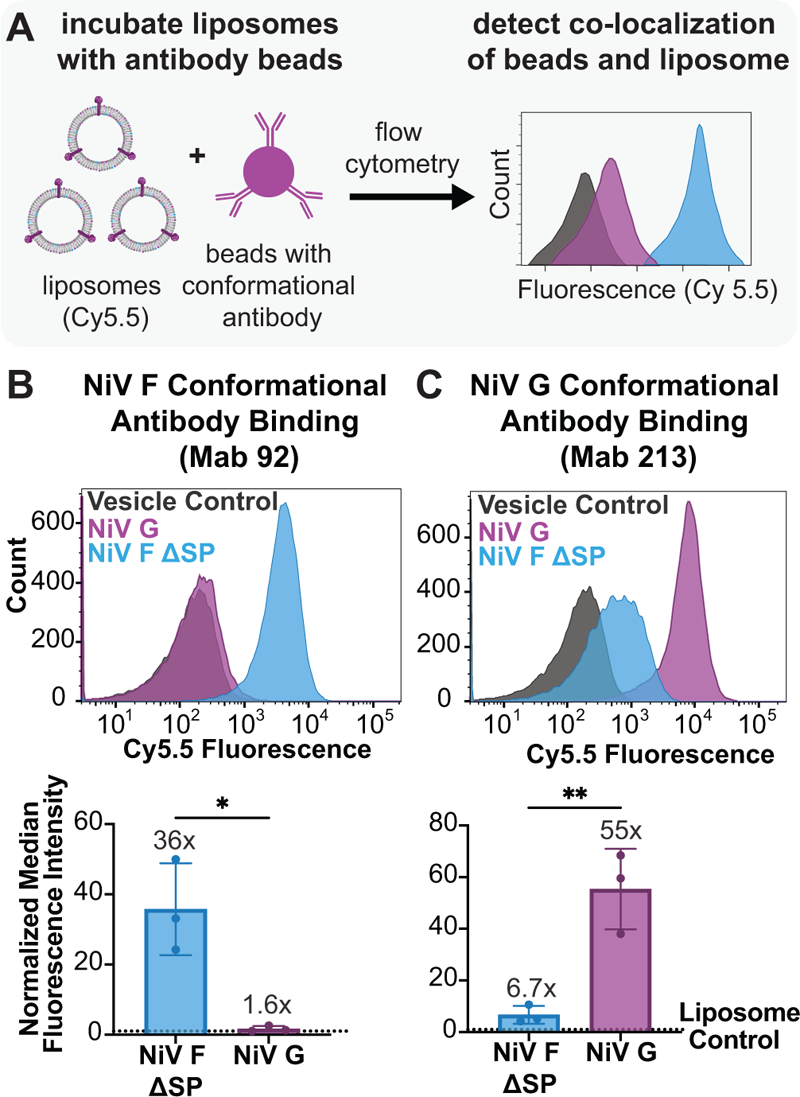
Cell-free expressed viral membrane proteins maintain native folding conformation. **(A)** Flow cytometry with conformational antibodies was used to detect native folding and orientation of NiV F ΔSP and NiV G upon expression and cotranslational integration into PC/PE/PS liposomes containing Cy5.5. **(B)** NiV F ΔSP and NiV G expressed in fluorescent PC/PE/PS liposomes as well as an empty vesicle control were incubated with beads functionalized with a monoclonal antibody (Mab 92) that binds prefusion F in the third helical region (HR3) of the NiV F protein. PC/PE/PS liposomes containing NiV F ΔSP bind beads with Mab 92 with a ∼36-fold increase in median fluorescence intensity relative to the empty vesicle control which is significantly greater than the liposomes containing NiV G (1.6-fold increase). (**C**) Beads were labeled with Mab 213, which binds to an epitope on the head region of NiV G. PC/PE/PS liposomes containing NiV G bind beads with a 65-fold increase in median fluorescence intensity relative to the empty liposome control compared to a 6.7-fold increase for NiV F ΔSP-containing liposomes relative to the empty liposome control. Error bars represent the standard deviation for n = 3 independent replicates. \**P* ≤ 0.05; \*\**P* ≤ 0.01; *P* values are from two-tailed t-tests.

For confirmation of NiV F ΔSP folding, we used Mab 92, which detects the native conformation of prefusion NiV F by binding prefusion F in the third helical region (HR3) of the NiV F protein (Fig. 3B) (75, 78). Flow cytometry revealed a large population of liposomes containing NiV F ΔSP bind to Mab 92-conjugated beads compared to both the no protein control (liposome control) and the nonspecific protein control (NiV G), as visualized by the rightward shift of Cy5.5 fluorescence for beads incubated with the NiV F ΔSP sample. The median fluorescence intensities of NiV F ΔSP and NiV G samples were normalized to the median fluorescence intensity of the vesicle control to account for background fluorescence due to nonspecific vesicle adsorption to antibody beads. Here, NiV F ΔSP expressed in PC/PE/PS liposomes displayed a ∼36-fold increase in median fluorescence intensity compared to the vesicle control while the NiV G sample had only about a 1.3-fold increase. Next, a similar study was performed to validate proper folding of NiV G using Mab 213, which binds to an epitope on the head region of NiV G (76), to detect the native conformation of NiV G (Fig. 3C). Again, we found that the median fluorescence intensity of beads colocalized to fluorescent liposomes containing NiV G displayed a ∼55-fold increase compared to the vesicle control. In contrast, the NiV F ΔSP displayed a 6.7-fold increase compared to the control, unmodified liposome, and was significantly lower in fluorescence than Mab 213 beads incubated with NiV G-containing liposomes. To further confirm proper folding of both proteins, we used Mab 66 and Mab 26 which also detect the conformations of NiV F and NiV G, respectively, yet bind distinct epitopes from Mab 92 and Mab 213 (75, 76). In similar studies, we found that higher levels of protein in liposomes bound to their respective antibody beads compared to nonspecific and no protein controls (Fig. S6). Lastly, we verified that MPLA does not affect the folding of NiV G by using Mab 213 and comparing NiV F ΔSP and NiV G expressed in 0% MPLA and 0.04% MPLA with a no protein control (Fig. S7). Together, our results indicate NiV F ΔSP and NiV G can be cell-free expressed in their native conformation and orientation in PC/PE/PS liposomes.

### Liposomes With Cell-free Expressed NiV Membrane Proteins Elicit Neutralizing Antibody Responses in Mice

To evaluate the immune response of the liposomes containing membrane integrated NiV F ΔSP and NiV G, we administered immunizations to distinct sets of mice intramuscularly. We screened a variety of conditions to identify CFE conditions for vaccine scale up. Several adjustments were made from the conditions used for liposomal characterization studies such as vesicle size (200 nm), plasmid concentration per reaction (400 ng), temperature (33°C), and time (6 hours), as these conditions were identified to yield the highest protein yield in purified vesicles during scale up (Fig. S8). Liposomes were assembled from a PC/PE/PS composition with either NiV F ΔSP, NiV F ΔSP/G, or NiV F ΔSP/G + MPLA and were subsequently purified and concentrated to achieve a dose of 2.5 µg/50 uL of liposomes. The total amount of MPLA per vaccine dose for the NiV F ΔSP/G + MPLA group was 16.5 µg. As a negative control, empty PC/PE/PS liposomes were employed. Mice received 50 µl (2.5 µg) of the vaccine preparation with 50 µl of alum or PBS (NiV F ΔSP/G + MPLA group) as an adjuvant and were boosted on day 21 post-prime vaccination ((female, n=3), Fig. 4A). The protein dosage administered was determined using a Coomassie assay. We monitored clinical symptoms and mouse weight for 7 days post-prime vaccination to assess potential adverse effects of the vaccine. No weight loss was evident among any of the vaccinated groups (Fig. 4B).

**Fig. 4.**
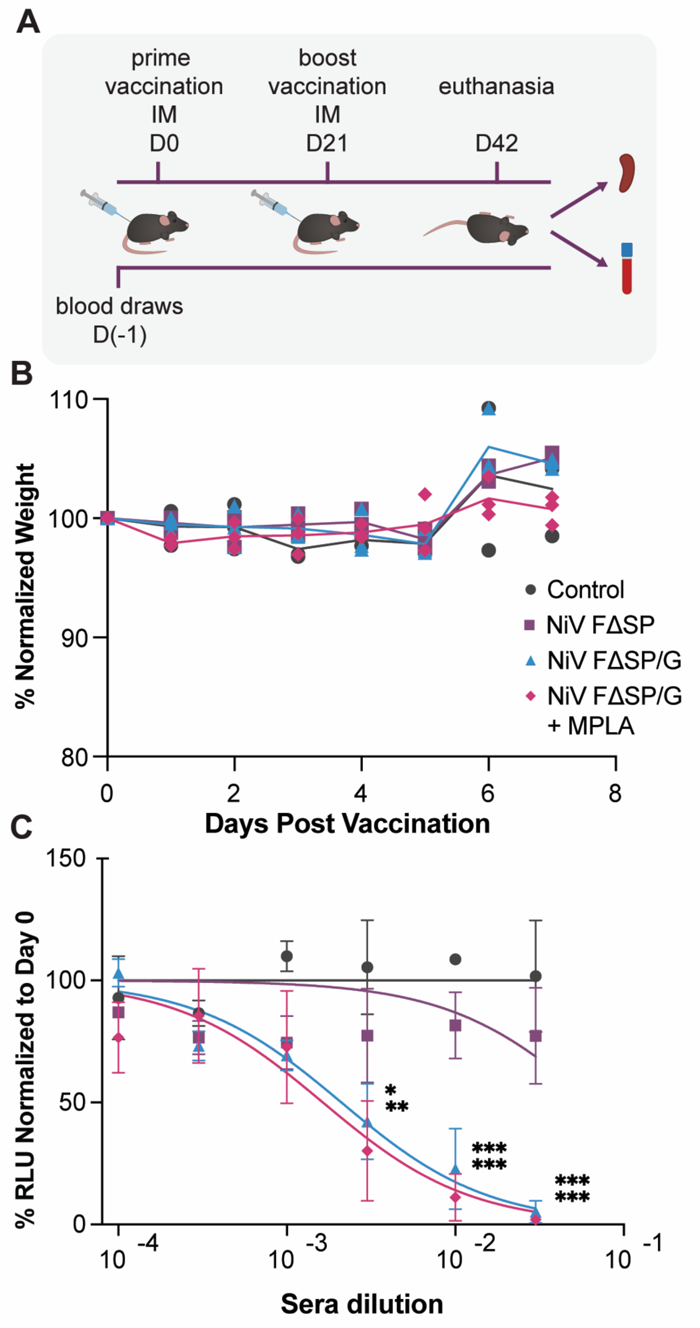
Liposomes containing cell-free expressed NiV membrane proteins induce humoral responses in mice (n=3). **(A)** A schematic illustrating the vaccination time points and sample collection. **(B)** Weight changes post-vaccination were monitored, and normalized to pre-vaccination weight on day 0. **(C)** The presence of neutralizing antibodies in mice sera were assessed by monitoring the neutralization of VSV-NiV F/G entry into Vero-E6 cells 42 days post-immunization. VSV-NiV F/G was treated with serial dilutions of sera from various groups of vaccinated mice, and the IC50 values were calculated. Significantly higher levels of neutralizing antibodies were observed in the NiV FG and NiV FG MPLA groups compared to the control group (***P value= 0.0002 and 0.0001 for a 1/30 dilution of sera, ***P value= 0.0009 and 0.0001 for a 1/100 dilution of sera, respectively, and *P value= 0.0192 for NiV FG and **P value= 0.004 for NiV FG MPLA for a 1/300 dilution of sera).Data is presented as mean values accompanied by the standard error of the mean (SEM) for n = 3. \**P* ≤ 0.05; \*\**P* ≤ 0.01; \*\*\**P* ≤ 0.001; *P* values are generated using two-way ANOVA and Tukey’s multiple comparisons test.

After vaccination, we assessed the immunogenic properties of our liposome vaccine platform in mice. Blood samples were collected weekly from day 0 to day 42. The presence of neutralizing antibodies in the final bleed serum was tested by assessing the capacity of this serum to inhibit infection of VeroE6 cells by luciferase gene-encoded pseudotyped vesicular stomatitis virus (VSV) incorporating NiV F/G. VSV infection was monitored with a Renilla Luciferase kit 24 hour post-infection. Fig. 4C presents the amounts of NiV F/G pseudotyped VSV virion neutralization with various dilutions of sera from the vaccinated mice. Mice vaccinated with the NiV F ΔSP/G and NiV F ΔSP/G + MPLA liposomes generated robust neutralizing antibody responses against NiV F/G VSVs. NiV F ΔSP/G and NiV F ΔSP/G + MPLA groups exhibited a half-maximal inhibitory concentration (IC50) at 1/500 and 1/600 dilutions of serum, respectively. The NiV F ΔSP/G + MPLA vaccinated group exhibited marginally elevated levels of neutralizing antibodies in their serum as compared to the NiV F ΔSP/G with alum group, indicating a potential enhancement in immune response attributed to the presence of the MPLA adjuvant. A greater difference in sera neutralization capacity was observed when comparing these vaccinated groups to NiV F ΔSP liposome vaccinated animals. This difference indicates that the neutralizing antibody responses to the multivalent F ΔSP/G vaccine were superior to the F alone vaccine, likely attributable to the heightened stimulatory effect of both F ΔSP and G proteins together compared to F ΔSP alone. As anticipated, no neutralizing antibodies were detected in the empty liposome vaccinated groups.

The T cell response to viral antigens post-vaccination was also assessed via *ex vivo* splenocyte stimulation. Splenocytes were collected from all vaccinated groups one month after the boost vaccination (Fig. 4A). These splenocytes were labeled with a dye, carboxyfluorescein diacetate succinimidyl ester (CFSE), the dilution of which is indicative of proliferation, then cultured for five days with media alone or pseudotyped NiV F/G VSV. Following culture, cells were harvested and subjected to flow cytometric analysis to quantify proliferated cells. The proliferation of CD4+ and CD8+ T cells (Fig. S9) were compared among the control, NiV F ΔSP, NiV F ΔSP/G, and NiV F ΔSP/G + MPLA vaccinated groups. The control and NiV F ΔSP vaccinated groups did not exhibit a significant change in T cell proliferation upon stimulation. In contrast, the T cells from both NiV F ΔSP/G and NiV F ΔSP/G + MPLA vaccinated groups exhibited higher, although not significant, proliferation upon stimulation with pseudotyped NiV F/G VSV, compared to unstimulated cells.

## Discussion

In this work, we present a cell-free expression platform for the assembly of multivalent liposomal vaccines that present properly folded, full-length viral membrane proteins. We targeted our platform to the Nipah virus by expressing two, full-length, viral membrane proteins: NiV F and NiV G. Critically, the viral membrane proteins folded into their expected, native conformations, and liposomes that displayed NiV F ΔSP and NiV G proteins and carried a lipid adjuvant elicited pathogen-specific IgG antibodies in mice.

Our approach is modular and highly tunable, allowing for changes in lipid carrier composition and the viral protein itself. With respect to lipid compositions, the range of lipids in this study was limited to changing glycerophospholipid compositions since these lipids are abundant in the plasma membrane. Future studies could vary other liposomal properties like hydrophobic thickness of the lipids constituting the liposomes, evaluate other charged lipids such as PA, PG, and PI, or include other lipids present in virus sites of membrane fusion like PI lipids and sphingomyelin. Formulations including other lipid adjuvants such as Pam3CSK4 or α-GalCer may also improve vaccine efficacy (36). Lastly, we included PS in the liposomal formulation for our study because the lipid improved CFE protein expression and integration for the NiV F protein. PS is an inner leaflet lipid that is externalized to the surface of apoptopic cells, acting as a signal for phagocytes to remove the cells (79). While this lipid can have immunosuppressive capabilities (80, 81), studies have demonstrated that exploitation of PS as a targeting moiety for antigen presenting cells can improve antigen uptake, antigen presentation, and antigen-specific antibody production (79, 82, 83). Future work should explore the impact of including these lipids and antigens on vaccine efficacy.

With respect to the viral proteins of interest, we focused our efforts on key membrane proteins in NiV, however, it should be possible to express a range of other viral membrane proteins that either cotranslationally integrate into membrane-based nanoparticles or colocalize to nanoparticles in other formats. In the current study, cotranslational integration of the NiV proteins into liposomes appeared to result in proteins that were oriented similarly to those in the native virus. However, an alternate orientation of a membrane protein may not necessarily limit vaccine efficacy. In fact, previous work has investigated antigen reorientation as a potential strategy for more broadly protective vaccines by directing immune responses to more conserved epitopes such as in the stem rather than the more variable head region of proteins (84). While we retained the transmembrane domain of proteins in this study as it has been found to be important for membrane protein conformation, it should also be possible to conjugate the soluble portion of viral membrane proteins onto liposomes using cell-free conjugation technologies like those we have previously demonstrated and should be explored in future work (85).

A pleasantly surprising result in our study was that the NiV F ΔSP and NiV G proteins, produced through a bacterial extract, bound conformational antibodies against forms of the protein produced in a mammalian host. Mammalian cells glycosylate proteins and glycosylation of viral membrane proteins is typically required for the proper folding, stability, and immunogenicity. However, glycosylation either requires proteins or cell-free extracts that are produced in mammalian cells, both of which are currently expensive and time consuming methods (50). In this study, we used the more available, lower cost, bacterial cell-free systems which do not contain machinery for glycosylation. While native viruses contain glycosylated proteins, it has been found in the context of other viral diseases that glycosylation is not necessary for influenza vaccine efficacy (86), and removal of glycans in some cases can expose conserved sequences, like with the SARS-CoV-2 spike protein, to provide broader protection against variants of concern (87). Preliminary experiments in mice evaluating response to vaccination showed a robust and encouraging antibody response, though a more subtle change in the T cell proliferation response. Our findings suggest glycosylation may not be necessary to elicit a robust immune response with the proteins expressed in this study and in fact, the absence of glycosylation alongside a properly folded membrane protein may expose more epitopes for antibody generation.

In summary, we present a platform that uses cell-free expression and cotranslational insertion of viral membrane proteins into liposomes to create a new vaccine candidate. Furthermore, we introduce important design parameters for vaccine assembly using cell-free integration of viral membrane proteins in liposomes, including liposome composition and protein sequence. We envision screening different membrane protein antigens, lipids, and adjuvant formulations for not only distinct pathogenic threats, but also toward universal vaccines that target multiple pathogenic threats. Accordingly, this platform should be broadly applicable to the rapid generation of multivalent vaccines against a wide variety of infectious diseases.

## Author Contributions

V.T.H., N.P.K., S.D., and H.C.A. conceived the project. H.C.A provided advice and reagents for the conformational antibody studies. V.T.H. performed and analyzed vaccine design and assembly characterization. E.S. performed, optimized, and analyzed scaleup and vaccine preparation for mice studies. S.E. helped design, performed, and analyzed all mouse vaccination studies. T cell proliferation studies were designed and performed by J.C., J.S., R.A. and analyzed by J.C., J.S., R.A., and A.A. V.T.H., N.P.K., and S.E. wrote the initial manuscript. All authors provided feedback and edits.

## Acknowledgments

The authors thank the Kamat lab for providing feedback for this project. We also thank Prof. James Moon for guidance on working with the lipid antigen, MPLA. This work was supported in part by the National Science Foundation under Grant No. 1935356 (S.D. and N.P.K.), Grant No. DMR-2145050 (N.P.K.), Grant No. 2021900 (V.T.H.), and the McCormick Research Catalyst Program (NPK). The work was also supported by NIH grant R35ES028244 (to A.A. and Gary Perdew), NIH R01 grant AI109022 to H.C.A. This work used the Northwestern RHLCCC Flow Cytometry Facility (NCI CA060553) and the Keck-II facility of Northwestern University’s NUANCE Center, which has received support from the SHyNE Resource (NSF ECCS-2025633), the IIN, and Northwestern’s MRSEC program (NSF DMR-2308691).

## Notes/Conflict of Interest Statement

V.T.H., N.P.K., S.D., E.S., S.E, and H.C.A. are inventors on a PCT application (PCT/US2023/076919) submitted by Northwestern University covering methods for assembling protein-conjugated nanocarrier vaccines.

## Materials and Methods

### Materials

1,2-dioleoyl-sn-glycero-3-phosphocholine (DOPC), 1,2-dioleoyl-sn-glycero-3-phosphoethanolamine (DOPE), 1,2-dioleoyl-sn-glycero-3-phospho-L-serine (sodium salt) (DOPS), 1,2-dioleoyl-sn-glycero-3-phosphoethanolamine-N-(cap biotinyl) (sodium salt) (18:1 Biotinyl Cap PE), and 1,2-dioleoyl-sn-glycero-3-phosphoethanolamine-N-(Cyanine 5.5) (18:1 Cy5.5 PE) were purchased from Avanti Polar Lipids. PURExpress® In Vitro Protein Synthesis Kit and PURExpress® Disulfide Bond Enhancer were from New England Biolabs. Phosphate-buffered saline (PBS) tablets were purchased from Sigma-Aldrich. Antibodies used include anti-FLAG (Abcam, F-tag-01), anti-Myc (Abcam, 9E10), anti-FLAG PE (Abcam, ab72469), and anti-Myc PE (Cell Signaling Technology, #3739), anti-mouse IgG HRP-linked (Cell Signaling Technology #7076). Conformational antibodies Mab 213, Mab 26, Mab 92, and Mab 66 were obtained as previously reported (74, 75). Pierce™ Streptavidin Magnetic Beads and Pierce™ Protein A/G Magnetic Beads were purchased from Thermo Fisher. 12% Mini-PROTEAN® TGX™ Precast Protein Gels, Immun-Blot® PVDF Membrane, and were purchased from Bio-Rad. NiV F construct was ordered from Twist Biosciences. NiV G construct was gifted from Susan Daniel’s lab. Primers were from IDT and cloning reagents were from Thermo Fisher (Phusion™ High-Fidelity DNA Polymerase, T4 DNA ligase) and New England Biolabs (T4 Polynucleotide Kinase).

### Methods

#### Protein Design and Gene Assembly

Protein sequences for NiV F and NiV G were obtained from UniProt (Q9IH63 and Q9IH62). Codon optimized sequences for expression in *E. coli* systems were designed with N-terminal FLAG (DYKDDDDK) and C-terminal Myc (EQKLISEEDL) tags. The constructs were ordered from Twist Biosciences in a pET21+ vector with a T7 promoter/terminator and added ribosome binding site. NiV F ΔSP with the signal peptide sequence (AA 1-26) deleted was generated using PCR amplification of the mutant and blunt-end cloning techniques.

#### Liposome Preparation

Liposomes were prepared via thin film hydration and extrusion. Lipid compositions in chloroform were mixed in a glass vial, dried with a stream of nitrogen gas, and placed in a vacuum for at least 3 hours to overnight. Films were rehydrated in Milli-Q water and incubated at 60°C for at least 1 hour. The rehydrated mixture was then briefly vortexed and extruded 21 times though a 0.1 µm polycarbonate membrane.

#### Cell-free Protein Synthesis of Membrane Proteins

Cell-free expression of the viral membrane proteins was performed using PURExpress *In Vitro* Protein Synthesis Kit following the manufacturer’s protocol with user added volume adjusted to 30 µL. PURExpress components, PURExpress Disulfide Bond Enhancer, 10 mM of liposomes, and ∼1.4 nM of DNA plasmid were assembled and incubated at 30°C for 8 hours.

#### Biotinylated Vesicle Purification and Semi-quantitative western blot Analysis

For vesicle purification for semi-quantitative western blot analyses, liposomes included 0.1 mol% 18:1 Cy5.5 PE and 1 mol% Biotinyl Cap PE. 3.5 µL of cell-free reaction mixture was mixed with 0.1 mg of Pierce Streptavidin beads, incubated at room temperature for 1 hour, and washed at least twice until supernatant A280 reached <0.05. Resuspended beads bound to the fluorescent biotinylated liposomes were read on a SpectraMax i3x plate reader at excitation 677 nm/emission 707 nm. Resuspended mixtures were mixed with 4x Laemmli buffer and loaded into 12% Mini-PROTEAN® TGX™ Precast Protein Gels for protein separation at 110V for 1:10 hours or 90V for 2 hours. Bands were transferred onto a PVDF via wet transfer at 100V for 1:10 hours. Membranes were then blocked for 30 minutes at room temperature in 5% milk in TBST buffer (20 mM Tris, 150 mM NaCl, 0.1% Tween 20, pH 7.6) then incubated in primary antibody (anti-FLAG or anti-Myc, 1:1000 dilution in 5% milk TBST) overnight at 4°C. Membranes were washed 3 times in TBST buffer, incubated in secondary antibody (anti-IgG, 1:3000 dilution in 5% milk TBST) for one hour at room temperature, then washed 3 times in TBST buffer. Before imaging on an Azure c280, membranes were incubated in Clarity Western ECL Substrate for 3 minutes. Western blot images were quantified using Fiji and values were normalized to bead fluorescence.

#### Flow Cytometry

For flow cytometry studies, liposomes included 0.1 mol% 18:1 Cy5.5 PE and 1 mol% Biotinyl Cap PE. For each sample, 20 ng of Pierce Protein A/G was incubated with 0.75 µL of antibody in PBS for 1 hour at room temperature, and washed twice to remove excess antibody. Antibody-bound beads were then incubated with 4 µL of cell-free reaction mixture for 1 hour at room temperature. Samples were washed twice more with PBS before flow cytometry reading. Samples were analyzed using a BD LSRFortessa™ Cell Analyzer with a 637 nm excitation laser and 730/45 nm emission. Liposome samples bound to beads were gated for size (FSC-A vs. SSC-A), singlets (SSC-W vs. SSC-H), and PE (PE antibodies only), and 50,000 events were collected. Data was analyzed using FlowJo.

#### Cell Cultures

The cell lines employed in this investigation were obtained from ATCC and utilized before reaching their 20th passage. Prior to experimentation, the cells were screened to ensure absence of mycoplasma contamination. Human embryonic kidney (HEK) 293T cells and Vero cells were cultured in Dulbecco’s minimum essential medium supplemented with 10% fetal bovine serum (FBS) and 1% penicillin/streptomycin. This culture medium was consistently used to maintain the cells throughout the duration of the study.

#### Immunization of mice

The protocol for immunizing mice was conducted in accordance with the guidelines approved by the Institutional Animal Care and Use Committee (IACUC). Twelve five-week-old C57BL/6 female mice (Charles River Laboratories) were acclimatized in the transgenic mouse core facility, Cornell University, for one week before initiating the immunization experiments. Each treatment group consisted of three mice, which were immunized intramuscularly with 50 µl of the test sample mixed with 50 µl of alum or MPLA as follows: group one received injections of bald liposome controls, group two received NiV F ΔSP, group three received NiV F ΔSP/G, and group four received NiV F ΔSP/G and MPLA liposomes. The total protein content in the test vaccines was 2.5 µg, determined using the BCA method. Mice received a vaccine boost on day 21. Blood samples were collected on days 0, 7, 14, 21, 28, 35, and 42, and the mice were euthanized one-month post-boost to collect spleen samples.

#### Production of NiV F/G Pseudotyped Virions

Recombinant NiV F/G VSV virions carrying a luciferase reporter gene were generated following the methodology previously described (88). Briefly, HEK 293T cells at 80–90% confluence were transfected with the pCAGGS expression plasmid encoding NiV F and G proteins for a duration of 16 hours using Lipofectamine 2000 (Invitrogen Inc.) in accordance with the manufacturer’s instructions. After transfection, the cells were exposed to a 1:30,000 dilution of recombinant VSVΔG containing the luciferase gene. The resulting pseudotyped VSV particles were then collected and purified for subsequent utilization in serum neutralization assays and ex vivo splenocyte activation studies.

#### Serum Neutralization

Serum samples obtained from vaccinated mice were subjected to dilution at ratios of 1:10, 1:30, 1:100, 1:300, 1:1000, 1:3000, and 1:10,000. The NiV F/G pseudotyped VSV was diluted at a ratio of 1:30,000. The pseudotyped VSV was treated with and equal volumes of each serum dilution. Following an incubation period of one hour at 37 °C, 100 µl of the mixture was added to Vero E6 cells at 40% confluency in duplicate and then incubated for 24 hours. Subsequently, the cells were lysed, and the mean luminescence activity was measured using the Renilla Luciferase assay following the manufacturer’s instructions (PromegaR)(1).

#### *Ex Vivo* Splenocyte Stimulation

Following euthanasia of mice at 1 month post-vaccination, spleens were collected, homogenized, and filtered using 70 micron filters following the methodology previously described (1). Briefly, ACK lysis buffer was utilized to lyse red blood cells, and splenocytes were then labeled with 5 μM CFSE according to the manufacturer’s instructions (cat# C34554; Thermo Fisher Scientific). Subsequently, the cells were plated into 24-well plates in media containing RPMI 1640 medium supplemented with 10% FBS, 4 mM L-glutamine, 0.1 mM nonessential amino acids, 1 mM sodium pyruvate, and 100 U/ml penicillin and streptomycin. The cells were either left untreated or exposed to the NiV F/G VSV pseudovirus for a duration of 5 days. Following incubation, the cultured mouse splenocytes were stained simultaneously with anti CD16/CD32 (Invitrogen), anti-TCR-beta-APC-cy7 (Biolegend), anti-CD4-Brilliant Violet 650 (Biolegend), and anti-CD8-Alexa Fluor 700 (Thermo Fisher) antibodies along with eBioscience Fixable Viability Dye-efluor-506 (Invitrogen). After washing, the cells were analyzed using the Thermo Fisher Attune NxT flow cytometer, with data analysis performed using FlowJo Software. Cell count data were normalized to unstimulated conditions of each mouse.

#### Liposomal Vaccine Preparation

All reagents and glassware used for vaccine preparation were sterilized by autoclaving including water, buffers, autoclave rotor, Eppendorf tubes, and pipette tips. Extruder equipment was cleaned using ethanol. Autoclave tubes and Amicon filters were only opened and loaded in a biological hood with proper ventilation. Liposome preparation, cell free reaction assembly, and sucrose gradient sample preparation for ultra-centrifugation were all completed in a biological hood under sterile conditions to minimize exposure to lipopolysaccharides. Unilamellar vesicles were produced using a combination of freeze/thaw cycles and extrusion. Briefly, lipids were combined in chloroform (mol ratio 15:4:1 of DOPC:DOPE:DOPS with 0.004 mol% MPLA), dried under a stream of nitrogen, and placed in a vacuum for a minimum of four hours. Lipids were then rehydrated in water and freeze/thawed. The vesicles were extruded 30 times through a 200 nm polycarbonate membrane. PUREfrex2.1 kits were used to synthesize the NiV F ΔSP and NiV G proteins according to manufacturer’s protocol with slight adjustments for optimized yield. Unless otherwise stated, the reagents included 10 mM liposomes, 1 mM cysteine, 2 mM DTT, ∼4.7 nM DNA plasmid, and PUREfrex2.1 solutions I, II, III. We found that adding DTT improved yield, likely due to preventing the formation of nonspecific disulfide bonds between the NiV F ΔSP protein subunits. The reagents were combined and incubated at 33°C for 6 hours unless otherwise stated.

#### Liposomal Vaccine Purification

An underlay step sucrose gradient method was used to separate the liposomes from the reaction components. A sterile 80% (w/v) sucrose solution was prepared in PBS. The top layer was 7 mLs of PBS (0% sucrose), the middle layer was 14 mLs of 25% (w/v) sucrose in PBS, and the bottom layer was ∼7 mL of 66% (w/v) sucrose. To prepare the bottom layer typically 5.87 mL of 80% (w/v) sucrose was combined with 1.2 mL of PUREfrex2.1 reaction mixture. A SW28 swinging bucket rotor was used to centrifuge the samples at 28,000 rpm for a minimum of 4 hours (8 for maximum recovery) at 4°C. The liposomes were recovered from fractions collected at the interface between PBS and 25 %(w/v) sucrose layers. Amicon spin filters 100kDa were used to concentrate the samples and remove sucrose. A minimum of three samples volumes were used to exchange the sucrose containing buffer into sterile PBS. The same procedures were followed for both the first and second mouse injections, but new reagents were prepared to maintain sterile conditions.

#### Liposomal Vaccine Protein Quantification

A combination of SDS-PAGE and a Bradford assay were used to quantify and evaluate sample purity of the proteins incorporated into the liposomes. Unless otherwise stated, the procedures for liposomal vaccine preparation and purification were used to assess the amount of protein incorporated into the particles. A minimum of a 4x PUREfrex reaction was required to be within the limit of the detection for SDS-PAGE analysis. Since the high concentrations of lipids are not compatible with most protein quantification techniques and distort the SDS-PAGE gels resulting in high background signal and smeared bands, the proteins were first removed from the vesicles using acetone precipitation. Acetone was cooled to -20°C and added in 4x volume excess to purified liposome sample. The solution was vortexed and incubated at -20°C for one hour. This mixture was centrifuged for 10 minutes at 13,000 x g and the supernatant was subsequently removed. The proteins were collected at the bottom as a pellet (typically not readily seen). To remove any remaining acetone, the tube was left open in a chemical hood to allow for any remaining acetone to evaporate for an additional 30 minutes. The protein was then dissolved in PBS buffer and analyzed against bovine serum albumin standard. Earlier experiments indicated the presence of higher-order structures, likely resulting from characteristic dimerization and trimerization of NiV F. To minimize the presence of these species, we added 5 mM DTT reducing agent to this solution and incubated for 1.5 hours at room temperature prior to analysis. It is likely that the protein yield was slightly underestimated due to potential product loss during the acetone precipitation. Nevertheless, the data shown in Fig. S8 indicate that these protocols resulted in a detectable and quantifiable protein yield with minimal unwanted contaminants. The SDS-PAGE analysis was run on 8% or 12% Bis-Tris PAGE gels in MOPS buffer at 175V for 1 hour (room temperature), and subsequently stained with Coomassie dye.

#### Liposomal Vaccine Protein Optimization

To maximize protein yield in the liposomes for vaccination studies, the reaction conditions were modified from those used for flow cytometry. To determine the best conditions a semi-quantitative western blot method was used to determine relative yields. The evolution of changing parameters is indicated in Fig. S8. Data shown in Fig. S8, A-D was collected as per the protocols outlined for liposomal vaccine preparation, purification, and quantification. For data shown in S.I. Fig 4 E-H, the methods were followed for liposomal vaccine preparation, but size-exclusion chromatography (SEC) was used to separate the vesicles from free proteins and the remaining PUREfrex components. This method was sufficient for optimization, however, was not used for vaccine purification as it resulted in lower purity. After samples were SEC purified, they were concentrated using 100kDa Amicon centrifugation filters and loaded on 8% or 12% PAGE gels. The same western blot procedure was followed as described in previous sections.

