## Supplementary Material for "Cell-free expression of Nipah virus transmembrane proteins for proteoliposome vaccine design"

**This PDF file includes:**

Figures S1 to S9
Tables S1
SI References

### Supplementary Figures

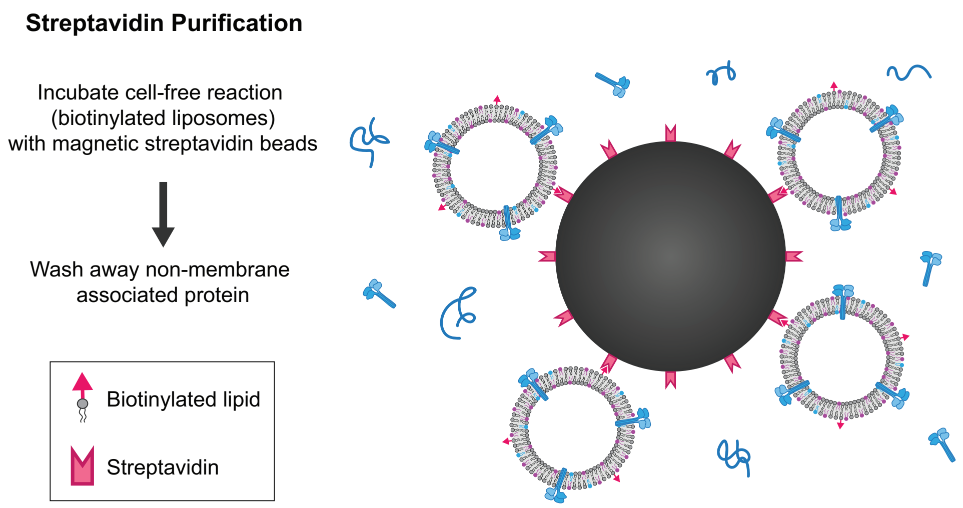

**Figure S1.** Proteoliposome purification scheme for semi-quantitative western blot analysis. After cell-free expression of proteins into liposomes containing 1% biotinyl cap PE lipid, samples are incubated with streptavidin beads to purify membrane associated proteins from aggregated and soluble protein in solution. Magnetic beads are washed at least two times following incubation before semi-quantitative western blot analysis.

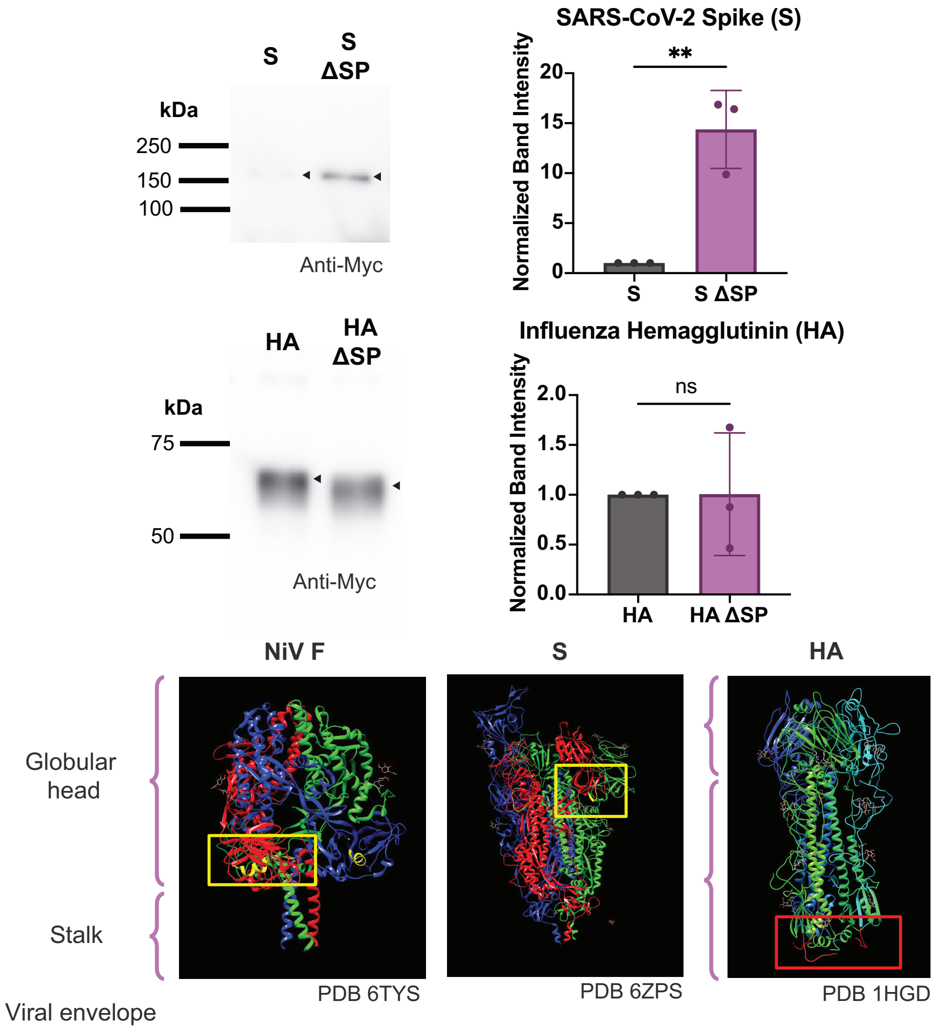

**Figure S2.** Signal peptides were removed (ΔSP) from the genes encoding SARS-CoV-2 Spike (S) and Influenza Hemagglutinin (HA). Proteins were expressed in PC liposomes then purified for semi-quantitative western blots. Antibody for the C-terminal Myc tag was used. Error bars represent the standard deviation for n = 3 independent replicates. $^{**}P \leq 0.01$; *P* values are from two-tailed t-tests.

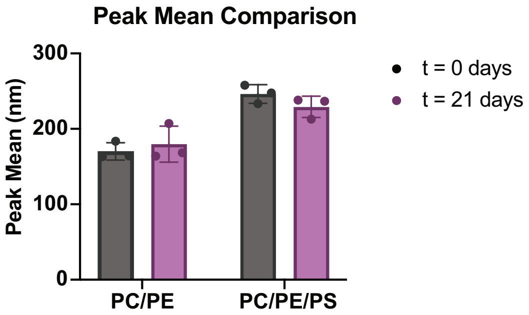

**Figure S3.** Comparison of the peak mean size of empty PC/PE and PC/PE/PS liposomes before and after 21 days (n = 3 measurements) using dynamic light scattering.

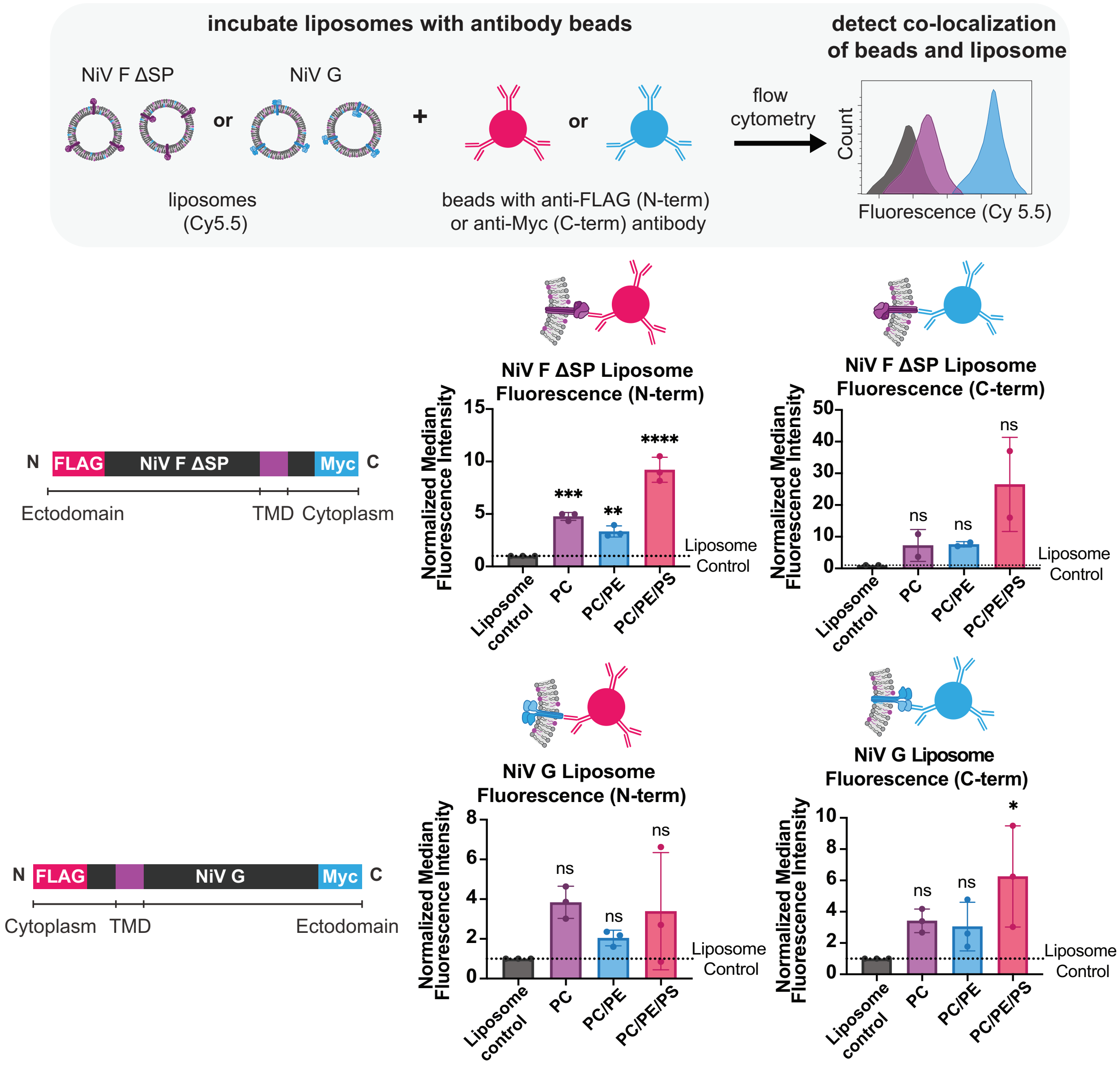

**Figure S4.** Flow cytometry was used to assess orientation of cell-free expressed NiV F ΔSP and NiV G. Protein A/G beads were labeled with PE conjugated anti-FLAG and anti-Myc antibodies that bind to the N-terminal FLAG tag and C-terminal Myc tags respectively. NiV F ΔSP or NiV G were expressed into PC, PC/PE, or PC/PE/PS liposomes containing 0.1% Cy5.5. Labeled beads were incubated with Cy5.5 fluorescent liposomes then run on the cytometer. Increased fluorescence indicates more protein oriented with the peptide tag outside of the liposome is binding to the bead. Median fluorescence intensity values of the beads for each liposome group were normalized to the median fluorescence intensity of the empty liposome control. Error bars represent the standard deviation for n=3, or n=2 independent replicates $^{*}P \leq 0.05$; $^{**}P \leq 0.01$; $^{***}P \leq 0.001$; $^{****}P \leq 0.0001$; $P$ values are generated using one-way analysis of variance (ANOVA) and Dunnett's multiple comparisons test for comparisons to the liposome control.

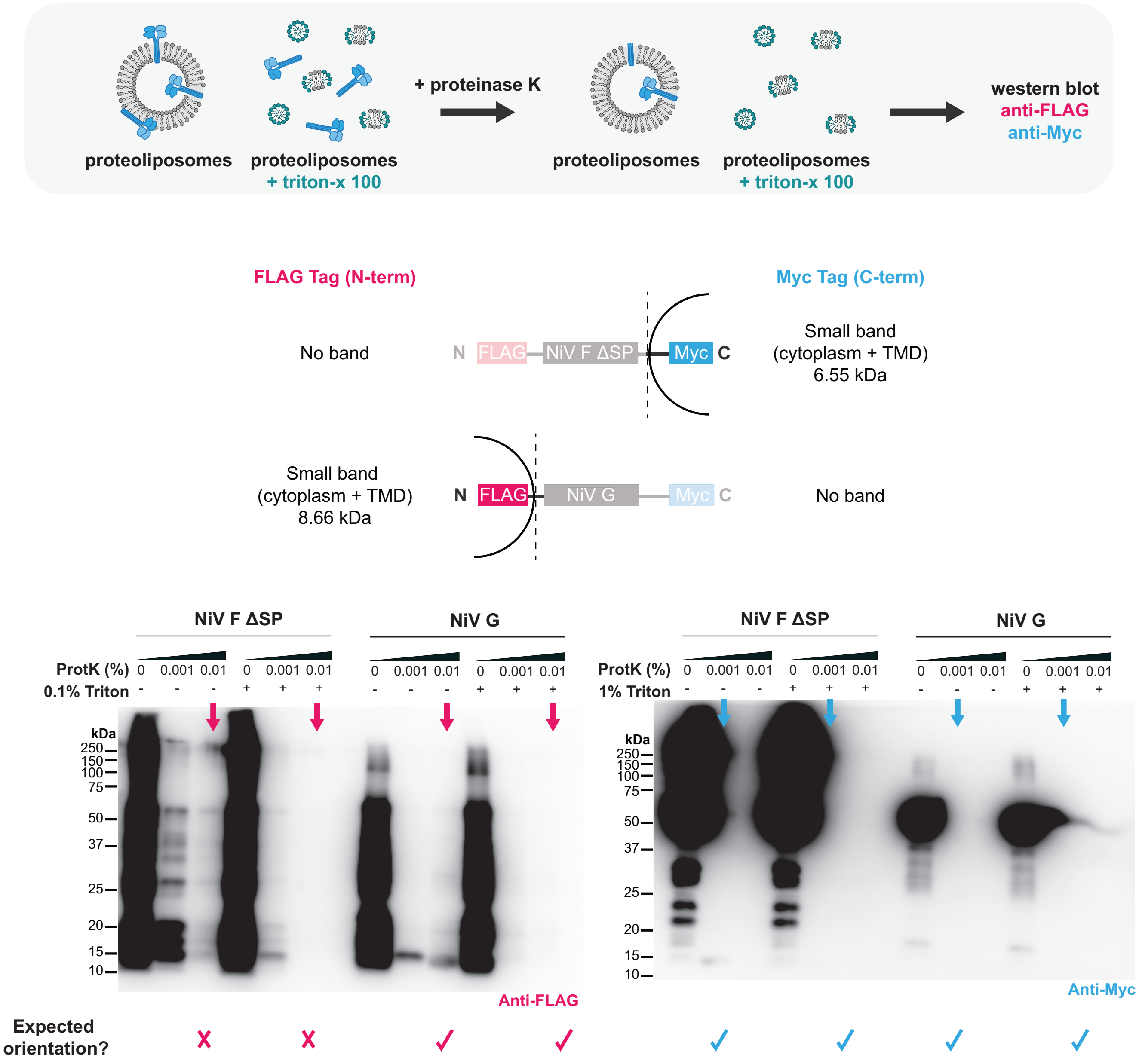

**Figure S5.** Proteinase K (ProtK) assay for assessing membrane protein integration vs. membrane association. Broad-spectrum protease ProtK digests protein outside of the liposome while protein integrated in the bilayer or in the liposome lumen should remain protected. NiV F ΔSP or NiV G were expressed in PC/PE/PS liposomes. Samples were incubated with increasing amounts of Proteinase K (0%, 0.001%, 0.01%). Samples had corresponding controls with 0.1% Triton X-100 to disrupt vesicles, indicating at what amount of ProtK there is total digestion of protein. Samples were then run on a western blot to visualize if N-terminal FLAG- and C-terminal Myc- tags are protected by the liposome, indicating integration and orientation of the protein. Samples with proteins integrated in expected orientation are denoted with a check mark whereas samples with proteins in the unexpected orientation are indicated with an X.

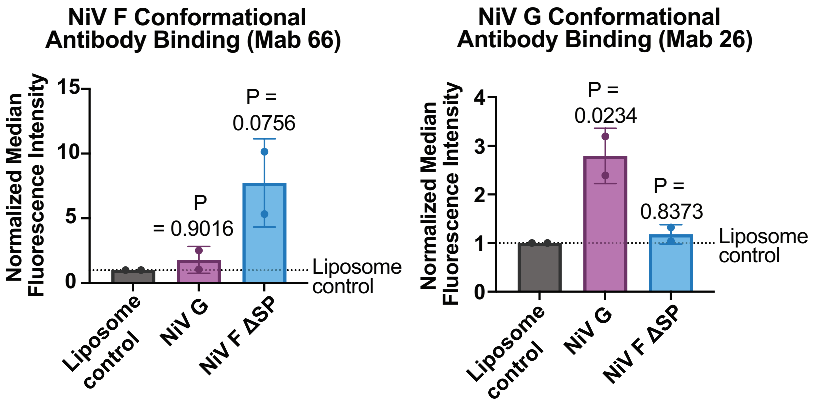

**Figure S6.** Two other conformational antibodies, Mab 66 for NiV F and Mab 26 for NiV G, were used in flow cytometry conformation studies to assess the proper folding if NiV F and G proteins, with a similar setup to Mabs 92 and 213 (1, 2). Error bars represent the standard deviation for n=2 independent replicates. *P* values are generated using one-way analysis of variance (ANOVA) and Dunnett's multiple comparisons test for comparisons to the liposome control.

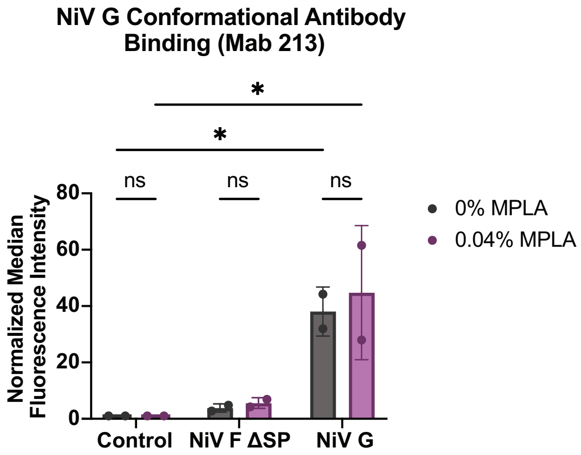

**Figure S7.** Conformational antibody Mab 213 for NiV G was used to compare protein folding of samples containing 0% or 0.04% MPLA using flow cytometry. In an assay similar to Figure 3 and Figure S6, NiV F ΔSP or NiV G were expressed in PC/PE/PS liposomes containing 0.1% Cy5.5 dye and 0% or 0.04% MPLA. Proteoliposomes were incubated with Protein A/G beads labeled with Mab 213, then beads were run on flow cytometry. Median fluorescence intensities of NiV F ΔSP or NiV G samples were normalized to respective empty liposome controls to determine a fold change. Error bars represent the standard deviation for n=2 independent replicates. *$P \leq 0.05$; $P$ values are generated using two-way analysis of variance (ANOVA) and Tukey's multiple comparisons test.

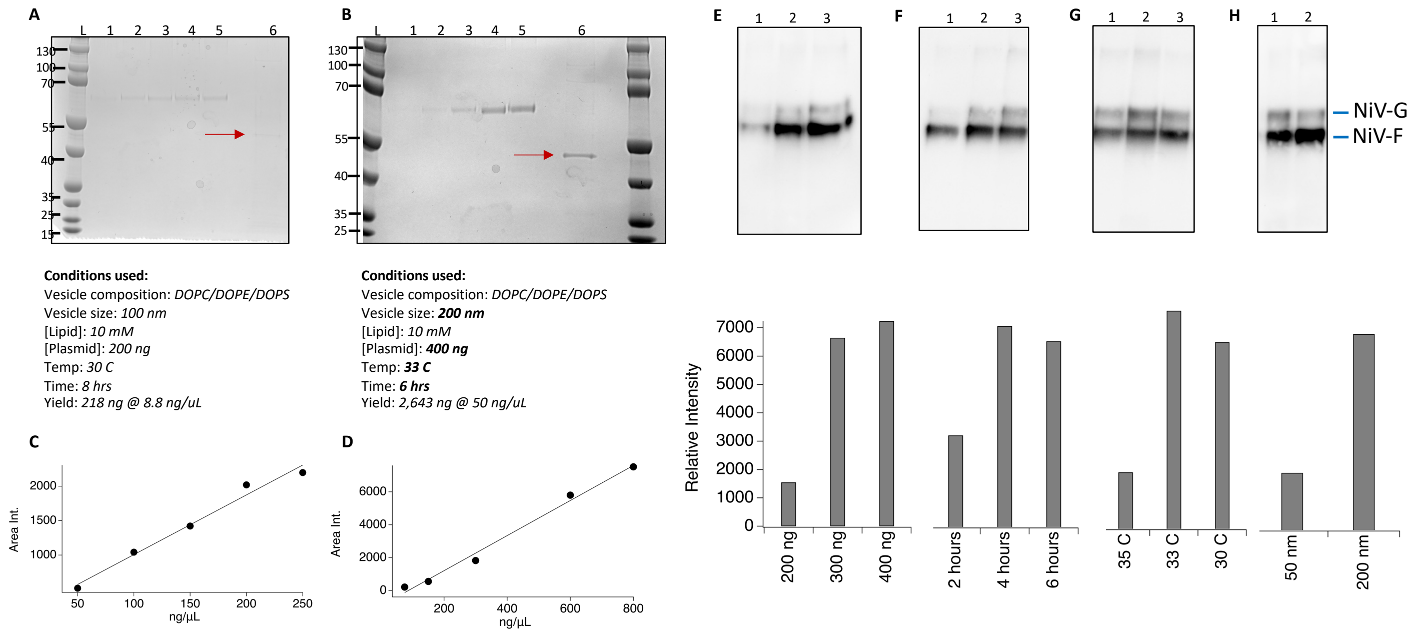

**Figure S8.** Quantification and optimization of Nipah proteins for scaled production of vaccine particles. A) SDS-PAGE gel of bovine serum albumin (BSA) standard in lanes 1-5 containing 50, 100, 150, 200, and 250 ng of proteins respectively. Lane 6 contains NiV F ΔSP proteins in 50 uL of purified liposomes. The conditions for this synthesis are indicated below the gel. B) SDS-PAGE gel of BSA standard in lanes 1-5 containing 75, 150, 300, 600, and 800 ng of proteins respectively. Lane 6 contains NiV F ΔSP proteins expressed with new conditions and are indicated below the gel. C/D) Calibration plots obtained from gels in (A) and (B) respectively using densitometry analysis using ImageJ and Igor64 software. The data points were fit to equation $Y = a + bx$ with $a = 138.4$, $b = 8.676$, and $r^2 = 0.98$ for data shown in (A), and $a = -916.39$, $b = 10.648$, and $r^2 = 0.99$ for data shown in (B). Using these values, the amount of NiV F ΔSP protein in the original conditions (A) was 35 ng and upon adjustment of parameters described in E-H was 423 ng for 50 uL of purified liposomes. The initial analysis of NiV F ΔSP shown in (A) revealed that although there was a sufficient concentration of proteins for flow cytometry, we were below the standard amounts required for injection into murine models and limited to a volume of <100 $\mu$Ls per mouse. To optimize [NiV F ΔSP] we tested variables such as E) [plasmid] in which lanes 1, 2, 3 show cell free reactions using 200, 300, and 400 ngs of initial NiV F ΔSP plasmids added to the cell free mixture; F) incubation times in which lanes 1, 2, 3 represent varying incubations periods of 2, 4, and 6 hours at 30 °C, where yield begins to decrease between 4-6 hours; G) temperature in which lanes 1, 2, 3 represent reactions completed at 35, 33, and 30 °C for 6 hours, and lastly H) vesicle size where lanes 1 and 2 had 50 and 200 nm vesicles added to the cell free reaction. Since all the other reactions tested contained 100 nm vesicles, we did not include that size on this western blot. All western blots shown in E-H used anti-FLAG monoclonal antibodies to detect the N-termini of the proteins. Though we appreciate that this method is semi-quantitative, we used these data to guide the improvement of proteins incorporated into the vaccine particles when scaled up, resulting in a 10-fold increase in [NiV F ΔSP]- which was sufficient for subsequent experiments completed with mice.

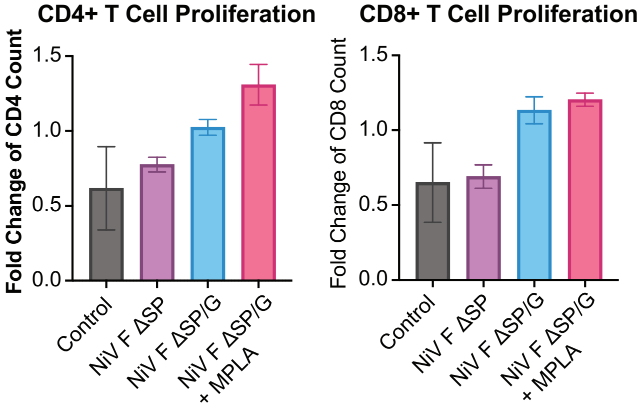

**Figure S9.** The *ex vivo* stimulation of splenocytes was monitored with a T cell proliferation assay (n=3). Cells were stained with the proliferation marker CFSE and stimulated with VSV-NiV F/G or media. On day 5 post-stimulation, staining with anti-TCRb, anti-CD4, and anti-CD8 antibodies to count these cell types. VSV-NiV F/G stimulated cells were normalized to unstimulated cells for each vaccination group.

**Supplementary table**

**SI Table 1.** Summary values of size and polydispersity (PDI) (n = 3) using dynamic light scattering.

| Liposome | Peak Intensity Mean (nm) (SD) | Polydispersity Index (SD) |
|---|---|---|
| NiV F ΔSP (PC) | 135 (23) | 0.21 (0.04) |
| NiV F ΔSP (PC/PE) | 128 (9) | 0.20 (0.02) |
| NiV F ΔSP (PC/PE/PS) | 110 (4) | 0.16 (0.02) |
| NiV G (PC) | 120 (8) | 0.17 (0.04) |
| NiV G (PC/PE) | 128 (30) | 0.17 (0.08) |
| NiV G (PC/PE/PS) | 134 (9) | 0.21 (0.02) |
| PC vesicle control | 123 (9) | 0.18 (0.05) |
| PC/PE vesicle control | 112 (4) | 0.18 (0.03) |
| PC/PE/PS vesicle control | 115 (8) | 0.20 (0.10) |
| PC vesicles (PBS) | 120 (6) | 0.18 (0.04) |
| PC/PE vesicles (PBS) | 129 (27) | 0.18 (0.06) |
| PC/PE/PS vesicles (PBS) | 121 (16) | 0.17 (0.04) |
